# Aggressive *KRAS* mutations direct TGF-β response towards partial EMT in patient-derived colorectal cancer tumoroids

**DOI:** 10.1101/2024.06.25.600620

**Authors:** Theresia Mair, Philip König, Milena Mijović, Loan Tran, Pedro Morata Saldaña, Carlos Uziel Pérez Malla, Kristina Draganić, Janette Pfneissl, Andreas Tiefenbacher, Julijan Kabiljo, Velina S. Atanasova, Jessica Kalla, Lisa Wozelka-Oltjan, Leonhard Müllauer, Michael Bergmann, Raheleh Sheibani-Tezerji, Gerda Egger

## Abstract

Transforming growth factor beta (TGF-β) exhibits complex and context-dependent cellular responses. While it mostly induces tumor-suppressive effects in early stages of tumorigenesis, its tumor promoting properties are evident in advanced disease. This TGF-β duality is still not fully understood, and whether TGF-β supports invasion and metastasis by influencing cancer cells directly, or rather through the stromal tumor compartment remains a matter of debate. Here, we utilized a library of colorectal cancer (CRC) patient-derived tumoroids (PDTs), representing a spectrum of tumor stages, to study cancer cell-specific responses to TGF-β. Using medium conditions allowing for the differentiation of PDTs, we observed TGF-β induced tumor-suppressive effects in early-stage tumoroids. PDTs with TGF-β pathway mutations or PDTs derived from metastatic tumors were insensitive to the treatment. Notably, one tumoroid line harboring an atypical *KRAS*^Q22K^ mutation underwent partial epithelial-to-mesenchymal transition (EMT), associated with morphological changes and increased invasiveness. On a molecular level, this was accompanied by elevated expression of mesenchymal genes, as well as deregulation of pathways associated with matrix remodeling and cell adhesion. Our results suggest that tumor cell intrinsic responses to TGF-β are critical in determining its tumor-suppressive or -promoting effects.

## Introduction

The TGF-β family of proteins includes various cytokines and growth factors, which are crucial for tissue integrity during development, homeostasis, and repair processes (1). TGF-β signaling affects multiple cell types and exhibits divergent roles either promoting or inhibiting cell proliferation, modulating cellular plasticity, remodeling the extracellular matrix (ECM), and regulating immune tolerance. Canonical TGF-β signaling is initiated by the activation of latent TGF-β and subsequent binding to hetero-dimeric TGF-β receptor complexes. This results in phosphorylation and activation of SMAD2/3 transcription factors (TFs), which subsequently bind to SMAD4. The SMAD-complex translocation to the nucleus eventually facilitates transcriptional activation of target genes in a context dependent manner through interaction with lineage-specific or signal-dependent TFs (1).

In healthy colon tissue, TGF-β modulates the differentiation and proliferation of epithelial cells and controls inflammatory responses to commensal bacteria, thereby maintaining tissue barrier functions (2). Perturbation of TGF-β signaling can promote inflammatory conditions such as Crohn’s disease or ulcerative colitis. Mutations in TGF-β pathway genes appear in 40-50% of colorectal cancers (CRCs) (3). Common mutations affect the *SMAD4* gene (10-35%) and the *TGFBR2* gene (15% in microsatellite-stable and more than 90% in microsatellite-instable CRCs). TGF-β also influences the tumor microenvironment by inducing gene expression changes in cancer-associated fibroblasts, thereby enhancing tumor aggressiveness, and by inducing immune suppression of tumor infiltrating cells of the innate and adaptive immune system (4,5).

Interestingly, while TGF-β acts as a tumor-suppressor in normal colon epithelia and early-stage colon cancer, it can induce EMT, which promotes tumor-progression and metastasis, in advanced stages (6,7). EMT, a plastic and multistep process, is characterized by morphological changes of cells, facilitating their detachment and motility (8). It was suggested that cancer cells retain some epithelial traits while they acquire mesenchymal characteristics, resulting in partial EMT states (9–12). The shift from tumor suppressive to tumor promoting functions is attributed to a decoupling of TGF-β signaling from apoptotic pathways or the activation of non-canonical pathways like PI3K-AKT or Ras-ERK-MAPK (13–16).

Advances in tumoroid models, which preserve molecular tumor characteristics, reflect tumor heterogeneity, and mimic patient response to therapies, have recently enhanced preclinical research (17). These models were also used to study TGF-β effects in physiologically relevant *in vitro* settings. For instance, murine-derived colon tumoroids with mutations in *Apc*, *Kras*, and *p53* genes exhibited partial EMT and collective invasion upon TGF-β treatment (18). Conversely, human CRC patient-derived tumoroids (PDTs) with active TGF-β pathway status mostly displayed tumor-suppressive effects, indicated by upregulation of CDK inhibitors, and in PDTs, which showed undisturbed proliferation, no signs of EMT were evident (4). Thus, the effect of TGF-β in CRC can be diverse and context dependent, and the mechanisms, which disconnect the tumor-promoting effects from its tumor-suppressive effects are still not fully understood.

In this study, we employed a series of 10 human CRC PDTs from tumors of different locations, progression stage, and mutational backgrounds to explore the dual nature of TGF-β effects in human CRC. PDTs with intact canonical TGF-β signaling predominantly exhibited tumor-suppressive and apoptotic responses to TGF-β. In contrast, lines with mutations in TGF-β pathway genes, or lines derived from CRC liver metastases, showed insensitivity to the treatment. Notably, one PDT line derived from a primary tumor with *APC* and atypical *KRAS* mutation, but with intact TGF-β signaling, displayed enhanced invasive properties, with phenotypic and molecular changes suggestive of partial EMT following TGF-β1 exposure. In summary, these results highlight the importance of the mutational background of CRC cells to promote tumor progression via cancer cell-intrinsic TGF-β responses.

## Results

### Generation of PDTs encompassing a spectrum of CRC progression

TGF-β elicits distinct responses in cancer cells depending on their state of progression and other contributing factors (19). Our study aimed to investigate this variability of TGF-β response in CRC PDTs. We utilized 10 different PDT lines derived from tumors with varying cancer grades (G2 or G3), and originating from different tumor locations, thereby covering a spectrum of CRC progression (**Fig. 1A**). Nine of the lines were *KRAS* mutant, while one MSI-high line harbored a *BRAF* mutation. During disease progression, CRC cells may develop resistance to TGF-β by acquiring mutations in key players of the canonical TGF-β signaling pathway, such as TGF-β receptors or SMAD proteins (3). Consequently, we grouped the PDTs generated from primary tumors into *SMAD4* wild-type (PDT1-PDT5) and *SMAD4* mutated (PDT6 and PDT7) lines. Moreover, the included MSI-high line carried a frameshift mutation in *TGFBR2* (PDT8), while two lines isolated from liver metastatic lesions had no mutations in TGF-β pathway genes (PDT9, PDT10). The included PDT lines did not only differ in their mutational background and origin, but also exhibited morphological diversity and predominantly mirrored the morphology of the original tumor tissue (**Fig. 1B**).

**Figure 1:**
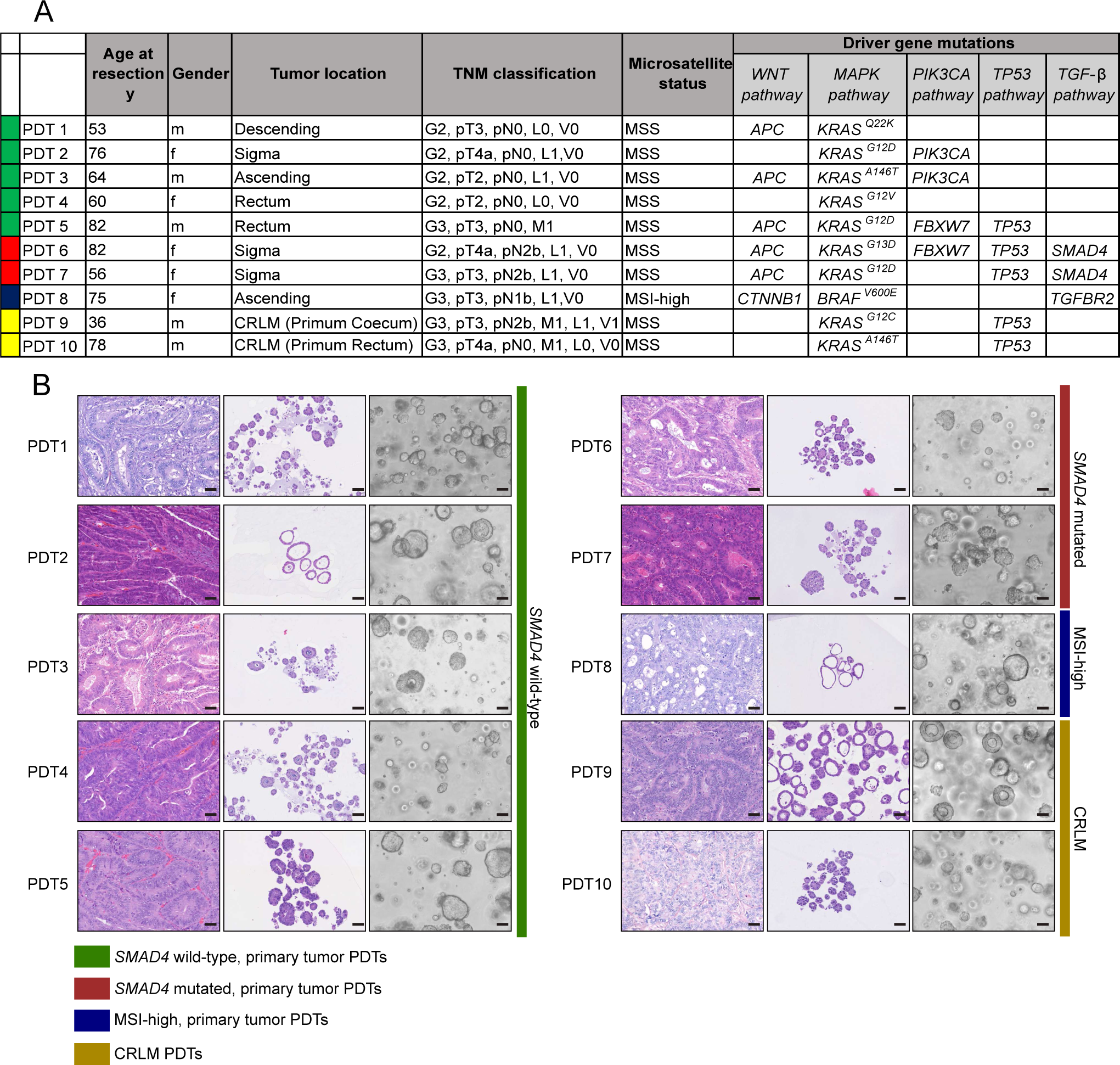
Molecular and phenotypic characteristics of CRC PDT lines. **A** Demographic table of different PDT lines used in this study, including patient age in years (y), gender, tumor location, TNM classification, microsatellite status, and driver gene mutations. The colors in the first row of the table group PDTs in four different classes. Green: *SMAD4* wild-type, primary tumor PDTs; red: *SMAD4* mutated, primary tumor PDTs; blue: MSI (microsatellite instable) high, primary tumor PDTs; yellow: colorectal liver metastasis (CRLM) PDTs. **B** Histo-morphological analysis of PDT lines compared to their tumors of origin. Hematoxylin and eosin (H&E) staining of FFPE sections of original tumor tissues (left) compared to PDT lines cultured for 7 days in ENAS medium (middle). Bright-field microscopic images of PDT lines cultivated for 7 days in ENAS medium (right). Scale bars: 50 µm. PDTs in B are grouped, and color coded as described in A.

### TGF-β1 treatment elicits divergent responses in CRC PDTs

To study the responses of the different lines to TGF-β1, the cell culture medium was adapted from standard organoid medium (ENAS, **supplementary table 1**), which had been developed for the growth of stem cells and contains TGF-β inhibitors among several other factors, to basal medium containing EGF as the only growth factor. After 10 days of culturing significant differences in the response of PDTs to the medium composition and TGF-β1 treatment were observed (**Fig. 2**). Some *SMAD4* wild-type lines including PDT2, 4 and 8 showed morphological changes upon culture in basal medium. For instance, PDT2 exhibited substantial morphological changes, transitioning from large, single-layered cystic PDTs to denser structures with thickened epithelium, indicating the differentiation of PDTs (20). As expected, long-term TGF-β1 treatment for 10 days had no suppressive effects on PDTs cultured in ENAS medium, due to the presence of TGF-β inhibitors in the medium. However, a tumor-suppressive effect of TGF-β1 was evident in most of the primary *SMAD4* wild-type PDTs in basal medium (PDT2-5), while lines carrying mutations in the TGF-β signaling pathway were resistant to TGF-β1 treatment (PDT6, 7, 8). Notably, PDTs derived from metastatic tumors (PDT9,10) showed only minor or no reduction in viability following TGF-β1 treatment in basal medium. Interestingly, the viability of PDT1, which was derived from a primary tumor with intact TGF-β signaling, was not affected by TGF-β1 administration. Additionally, the morphological changes, which were apparent in basal medium, were further enhanced in this line upon treatment with extended 2D growth patterns.

**Figure 2:**
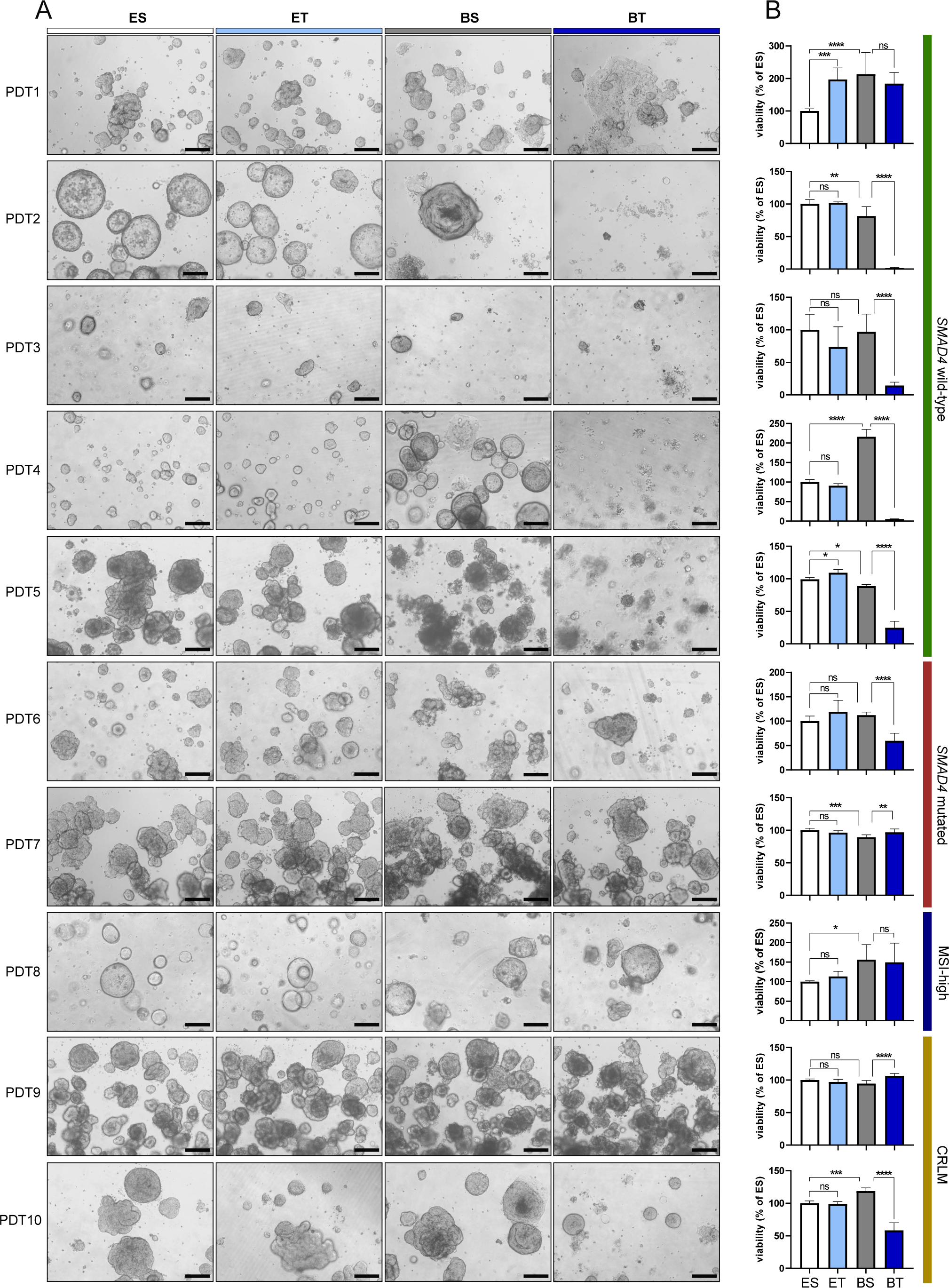
TGF-β1 treatment induces different responses in individual PDTs. **A** Representative bright-field microscopic images of different PDT lines cultured in different media conditions for 10 days, ES: ENAS + solvent (white); ET: ENAS + 5 ng/ml TGF-β1 (light blue), BS: basal medium + solvent (grey); BT: basal medium + 5 ng/ml TGF-β1 (dark blue). Scale bar: 200 µm. **B** Cell viability of PDTs in different media conditions as in (A) measured with CellTiter-Glo® 3D Cell Viability Assay. Viability is presented as % of viability relative to ES. Graphs show three technical replicates of two individual experiments (n=2). Statistical significance was calculated with GraphPad Prism version 8 using ordinary one-way ANOVA followed by Tukey’s multiple comparison test with 95% confidence interval. Ns P > 0.05; *P f 0.05; **P f 0.01; ***P f 0.001; ****P f 0.0001.

In order to investigate the different effects of TGF-β1 treatment on primary *SMAD4* wild-type lines in more detail, we assessed SMAD2/3 protein abundance and SMAD3 phosphorylation, as well as apoptotic signaling *via* the TGF-β-induced pro-apoptotic factor BIM (13,21) (**Fig. 3A, B**). PDT1, which did not show reduced viability following treatment, and PDT4, which showed reduced viability, were exposed to TGF-β1 for five days (**Fig. 3A**). Intriguingly, PDT1 displayed high levels of SMAD3 phosphorylation and modest amounts of BIM expression following treatment (**Fig. 3B**). Conversely, PDT4 exhibited strong SMAD3 activation in addition to increased BIM protein levels, suggesting increased pro-apoptotic signaling, elucidating the tumor-suppressive effects of the treatment. Furthermore, BIM protein levels were anti-correlated to phospho-ERK levels following TGF-β1 treatment in the PDT1 line. Activated ERK was previously shown to promote BIM phosphorylation resulting in its degradation (22). Furthermore, the expression of the pro-survival factor BCL-XL was induced in the PDT1 line upon TGF-β1 administration, additionally reinforcing the survival advantage of this line compared to PDT4, where no increased levels of BCL-XL were detectable. Similarly, 10-day TGF-β1 treatment of the primary PDT1 line, as well as the metastatic lines PDT9 and 10, displayed activation of SMAD3 and reduced BIM protein levels, which was paralleled by ERK activation in PDT1 and PDT10 (**Supplementary** Fig.1).

**Figure 3:**
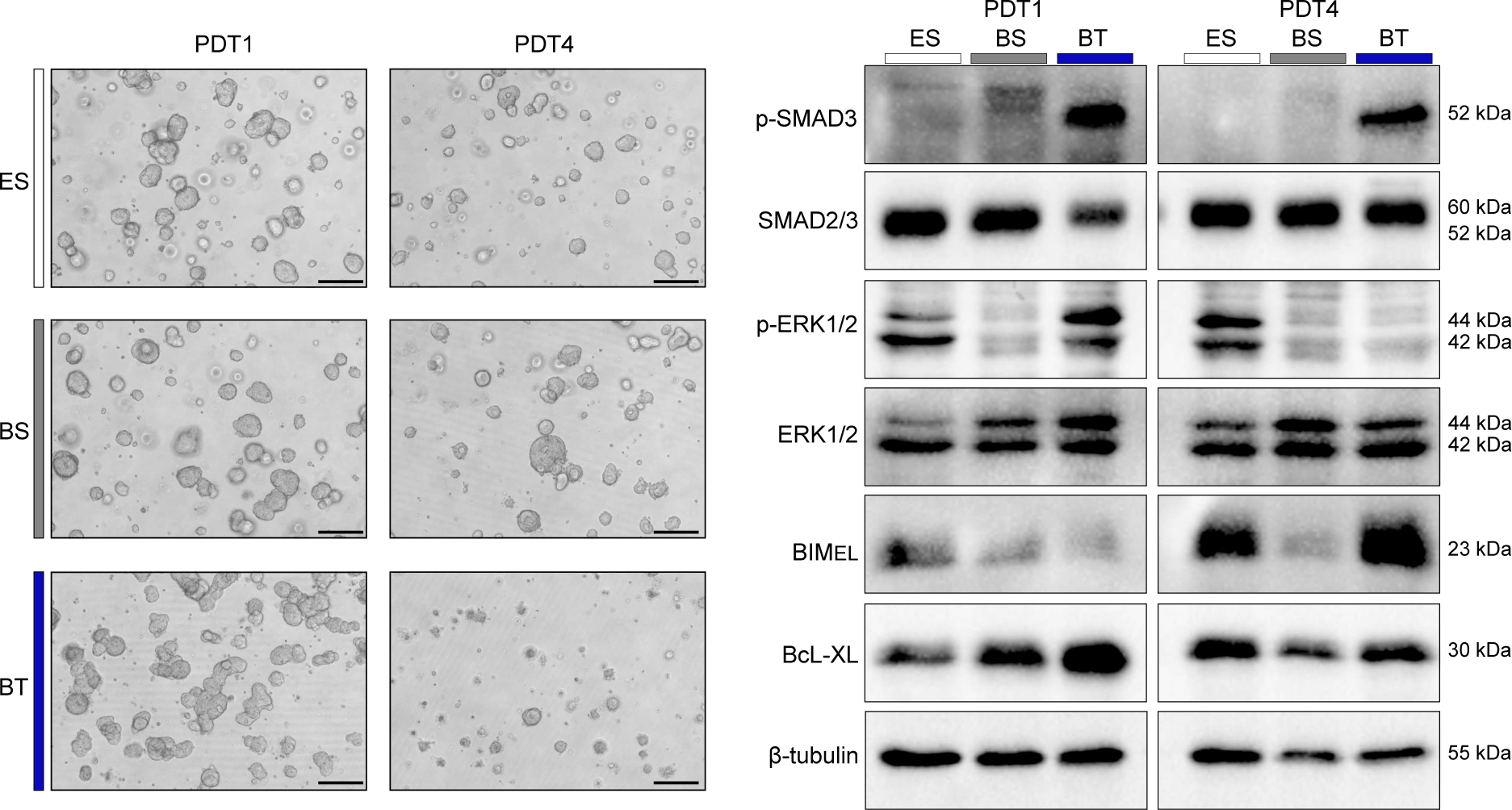
TGF-β1 induces different signaling pathways in insensitive and sensitive PDTs. **A** Representative bright-field microscopic images of PDT1 (TGF-β1 insensitive) and PDT4 (TGF-β1 sensitive) cultured in different conditions for five days. ES: ENAS + solvent; BS: basal medium + solvent; BT: basal medium + 5 ng/ml TGF-β1. Scale bar: 200 µm. **B** Western blot analysis of PDTs treated as in (A) for phospho-SMAD3 (p-SMAD3) (53kDa), total SMAD2/3 (60kDa, 52kDa), phospho-ERK1/2 (p-ERK1/2) (44kDa, 42kDa), total ERK1/2 (44kDa, 42kDa), BIM_EL_ (23kDa), BcL-XL (30kDa) and β-tubulin (55kDa) as loading control (n=2).

Together, these data indicate that TGF-β1 responses of PDTs largely reflect the diverse *in vivo* effects of TGF-β signaling. While tumor-suppressive consequences were observed in most of the primary *SMAD4* wild-type PDTs, metastatic and more advanced PDTs were affected less by the TGF-β1 treatment.

### PDT culture in basal medium induces differentiation towards specialized cell types of the colon

To further investigate the effects of TGF-β1 treatment on the primary PDT1 line, which showed no change in viability but strong morphological differences, we assessed the gene expression changes of this specific line under different medium conditions and following TGF-β1 treatment (**Fig. 4**). Principal component analysis (PCA) revealed clustering of PDTs cultivated in ENAS medium with or without TGF-β1, whereas PDTs cultivated in basal medium with or without TGF-β1 showed additional significant alterations and clustered separately (**Fig. 4A**). Unsupervised hierarchical clustering based on the top 1000 most variable expressed genes grouped PDTs cultured in ENAS medium apart from PDTs grown in basal medium +/- TGF-β1 (**Fig. 4B**). Similarly, differential gene expression analysis resulted in only 28 significantly deregulated genes (18 up, 10 down) (adjusted P-value (Padj) < 0.05. log2 fold change (LFC) > 1) in ENAS *versus* ENAS-TGF-β1 treated PDTs (**Fig. 4C**), while 3737 significantly deregulated genes (2197 up, 1540 down) were detected in the ENAS *versus* basal medium comparison (**Fig. 4D**). Moreover, a strong impact of TGF-β1 treatment was observed in basal medium, with 1570 significantly deregulated genes (647 up, 923 down) (**Fig. 4E**).

**Figure 4:**
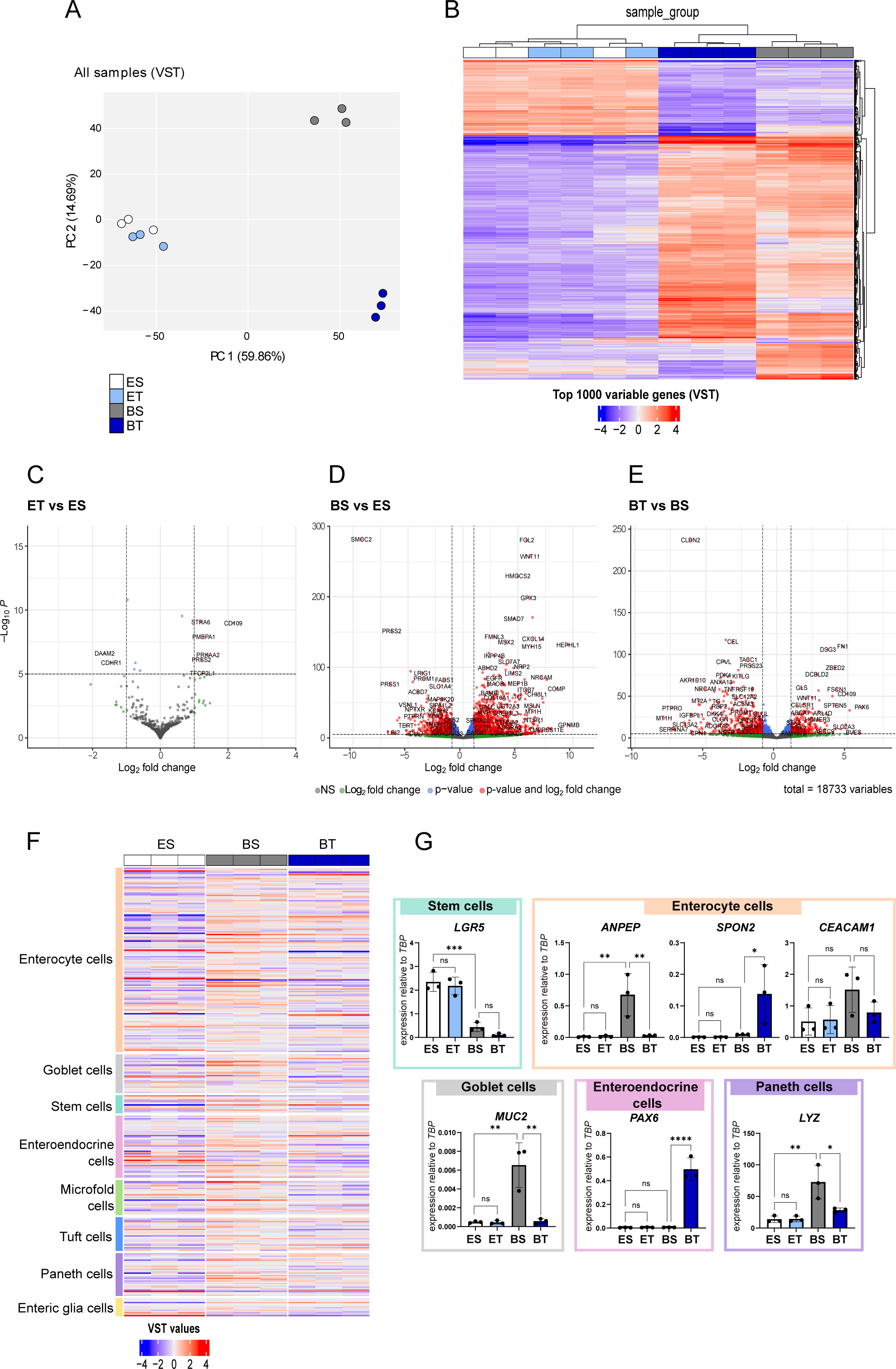
Gene expression analysis reveals differentiation of PDTs to specific cell types of the colon crypt in basal medium. **A** Principal component analysis (PCA) of RNA-sequencing data of PDT1 cultured in different conditions based on variance stabilizing transformation (VST) read counts (n=3). Colors indicate different conditions: white = ES: ENAS + solvent; light blue = ET: ENAS+ 5 ng/ml TGF-β1; grey= BS: basal medium + solvent; dark blue= BT: basal medium + 5 ng/ml TGF-β1. **B** Dendrogram and heatmap showing unsupervised hierarchical clustering of the top 1000 variable expressed genes using normalized read counts (VST transformed counts in DESeq2) of PDT1 cultured as in (A). Columns represent individual samples (color coded as in A), rows represent individual genes. The color gradient on the bottom shows VST normalized counts, with blue indicating below-average gene expression and red indicating above-average expression. **C-E** Volcano plots of significantly differentially expressed genes between different media conditions as in (A). Significantly deregulated genes are indicated in red with Padj < 0.05. LFC> 1. Genes below this significance threshold are indicated in grey (Padj > 0.05. LFC < 1), green (Padj > 0.05. LFC > 1) and blue (Padj < 0.05. LFC < 1). **F** Heatmap showing VST gene count values of PDT1 in different conditions (ES, BS, BT) for colon crypt cell type-associated genes inferred from (23) (https://panglaodb.se/). Cell types are ordered according to their frequency in the colon crypt. Red indicates upregulation, blue downregulation. **G** Gene expression analysis of indicated cell type-associated genes in different conditions (as in A) quantified by qRT-PCR. Data are represented as expression relative to TATA-Box Binding Protein (*TBP*) as housekeeping gene. Each dot represents the mean of 3 technical replicates (n=3). Statistical significance was calculated using GraphPad Prism version 8 with ordinary one-way ANOVA followed by Tukey’s multiple comparison test with 95% confidence interval: ns P > 0.05; *P f 0.05; **P f 0.01; ***P f 0.001; ****P f 0.0001.

Next, we explored gene expression levels of marker genes, which are characteristic for various cell types of the colon crypts (23). Notably, genes associated with enterocytes and secretory cells were generally upregulated in basal medium conditions compared to ENAS, whereas stem cell markers were downregulated (**Fig. 4F, supplementary Fig. 2A**). Upon addition of TGF-β1, the expression of most of these cell type-associated genes was dampened. Notably, some specific genes such as *SPON2*, a gene associated with enterocytes and CRC progression (24), *PAX6*, encoding a TF associated with enteroendocrine differentiation (25), and the goblet cell-associated gene *KRT7*, which has been linked to metastasis (26), were specifically upregulated in basal media with TGF-β1 (**Supplementary** Fig. 2). The expression of several genes was confirmed by qRT-PCR also highlighting the downregulation of the stem cell marker *LGR5* in basal medium conditions, and altered expression of enterocyte (*ANPEP*, *SPON2*, *CEACAM1*), goblet (*MUC2*), enteroendocrine (*PAX6*) and Paneth cell (*LYZ*) associated genes in the different conditions (**Fig. 4G**).

Taken together, these data demonstrate the upregulation of genes associated with different colon crypt cell populations upon withdrawal of stem cell niche factors and an additional deregulation of genes involved in tumor progression and metastasis upon TGF-β1 stimulation.

### TGF-β1 treatment of PDT1 in basal medium shifts the differentiation of PDTs towards a mesenchymal phenotype

To gain deeper insights into the molecular changes of the PDT1 line following TGF-β1 treatment, we used the CMS caller tool (27) to determine the consensus molecular subtypes (CMS), which have been used to classify CRCs in four different sub-types based on gene expression signatures (28). While PDTs grown in ENAS conditions could not be attributed to a specific CMS subtype, PDTs grown in basal medium were classified as CMS2 (canonical) or CMS3 (metabolic) (**Fig. 5A**). Notably, PDTs cultivated in basal medium with TGF-β1 were classified as CMS4, representing the mesenchymal sub-type of CRC.

**Figure 5:**
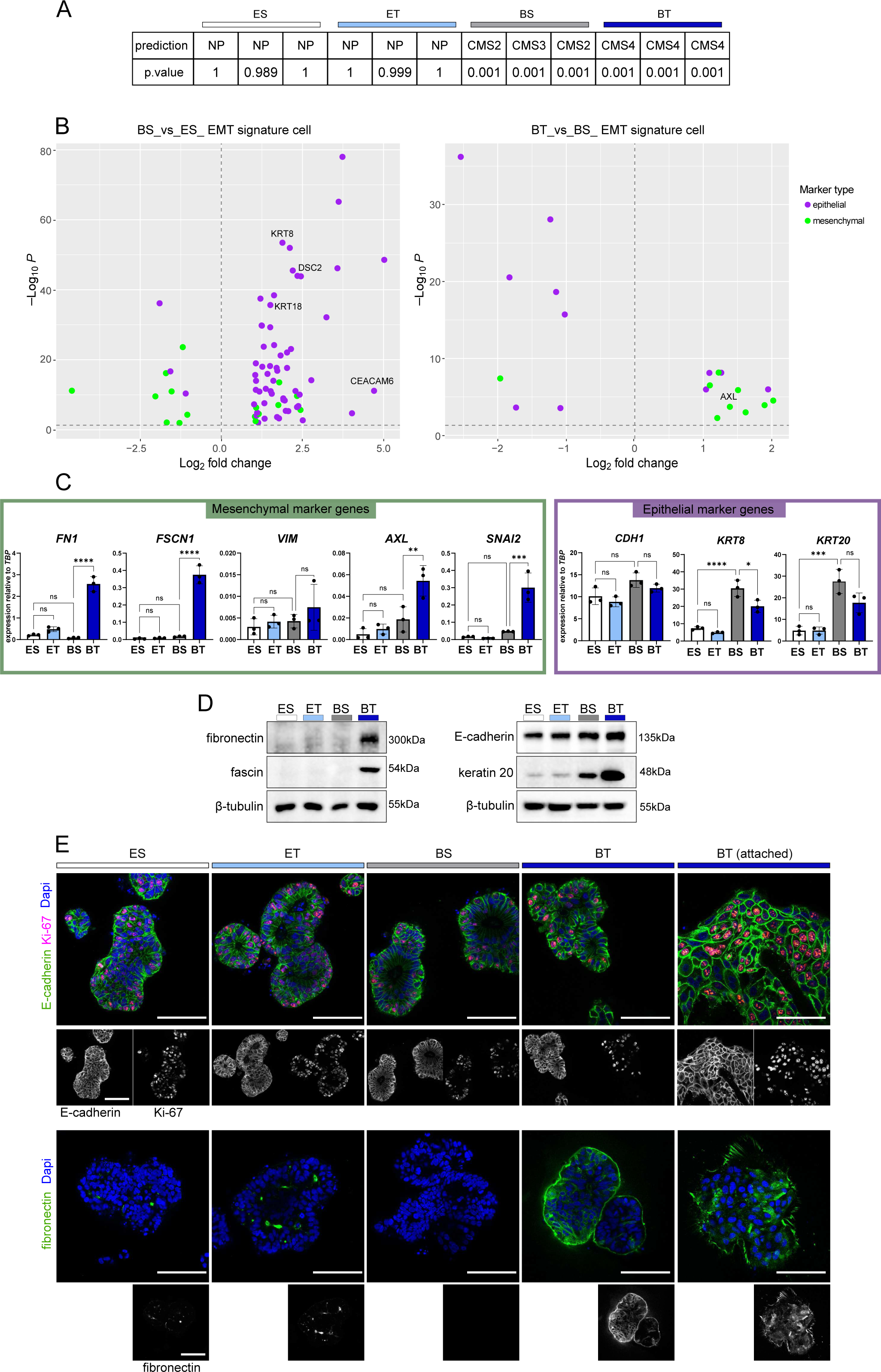
TGF-β1 treatment of PDT1 promotes differentiation towards a mesenchymal phenotype. **A** Prediction of the consensus molecular subtypes (CMS) of PDT1 cultured in different medium compositions (ES: ENAS + solvent; ET: ENAS + 5 ng/ml TGF-β1, BS: basal medium + solvent; BT: basal medium + 5 ng/ml TGF-β1) from RNA-seq data (n=3) using the CMS caller tool (27). The predicted CMS is indicated in the first row as NP: no prediction, CMS1, CMS2, CMS3 or CMS4 for each sample. The second row shows the P-values for each prediction. **B** Overlap of differentially expressed genes of PDT1 cultivated for 10 days in different conditions (BS vs ES and BT vs BS) with cancer cell specific EMT signatures (29). Volcano plot on the left represents BS *versus* ES, plot on the right represents BT *versus* BS. Epithelial genes are represented in purple and mesenchymal genes in green. **C** Gene expression analysis of mesenchymal genes (left) and epithelial genes (right) using qRT-PCR of PDT1 cultured in different conditions as in (A). Gene expression of target genes is represented as expression relative to *TBP* as housekeeping gene. Each dot represents the mean of 3 technical replicates (n=3). Statistical significance was calculated using GraphPad Prism version 8 with ordinary one-way ANOVA followed by Tukey’s multiple comparison test with 95% confidence interval: Ns P > 0.05; *P f 0.05; **P f 0.01; ***P f 0.001; ****P f 0.0001. **D** Representative Western blot analysis of selected mesenchymal (left) or epithelial (right) marker proteins isolated from PDT1 treated as in (A) (n=2). β-tubulin is used as loading control. **E** Whole-mount immunofluorescence staining and confocal microscopy of PDT1 cultured in different conditions as in (A) (n=2). The upper panel shows staining with antibodies against E-cadherin as epithelial marker (green) and Ki67 as a marker for proliferative cells (red). Nuclei were counterstained with DAPI (blue). The lower panel shows staining with antibodies against fibronectin as mesenchymal marker (green), and DAPI counterstain of nuclei (blue). Single channel pictures for each antibody are shown below individual multi-channel pictures in black and white. Scale bar: 100 µm.

Next, we compared gene expression profiles of this line in different media and after TGF-β1 treatment to published EMT signature genes (29). Cultivation of PDTs in basal medium induced the expression of epithelial marker genes, genes coding for intermediate filament-forming keratins such as *KRT8/18,* and cell adhesion molecules including *CEACAM1* and *DSC2* compared to the ENAS-cultured PDTs (**Fig. 5B, left**). Supplementation of basal medium with TGF-β1 induced a shift from epithelial to mesenchymal gene expression signatures (**Fig. 5B, right**). This was also confirmed by qRT-PCR with significant upregulation of fibronectin (*FN1*), fascin (*FSCN1*), and the receptor tyrosine kinase *AXL*, which has been associated with EMT, tumor cell invasion, and therapy resistance in different tumor entities (30) (**Fig 5C**). Moreover, the expression of the EMT transcription factor *SNAI2* was upregulated upon TGF-β1 treatment. For the mesenchymal gene vimentin (*VIM*), a trend of higher expression after TGF-β1 treatment was detectable. Epithelial marker genes were generally expressed at higher levels in basal medium compared to ENAS medium, confirming the differentiation phenotype of the stem cell-like cultures. However, the addition of TGF-β1 showed a trend towards lower expression of epithelial markers including E-cadherin (*CDH1*), *KRT8* and *KRT20* (**Fig. 5C**). Similarly, we detected a strong induction of the mesenchymal marker’s fibronectin and fascin on protein levels and high levels of the epithelial proteins E-cadherin and keratin 20 in all conditions (**Fig. 5D**).

Using immunofluorescence analysis, we evaluated the protein expression of E-cadherin and fibronectin under different conditions in the PDT1 line. Moreover, proliferation rates of PDTs were evaluated by Ki-67 staining, which showed similar proliferation rates of PDTs in different conditions (**Fig. 5E**). For PDTs grown in basal medium containing TGF-β1, we assessed both 3D structures and cells that had attached to the plates in 2D. E-cadherin expression was detected to similar levels in all conditions (ENAS, basal medium, with/without TGF-β1), whereas a strong induction of fibronectin was apparent only in PDTs cultured in basal medium after TGF-β1 treatment. Importantly, cells that grew in 2D showed prominent fibril formation, a key process in TGF-β induced EMT (31).

Collectively these results indicate that treatment with TGF-β1 shifts the differentiation of PDTs towards a mesenchymal phenotype, showing gene expression characteristics of partial EMT characterized by simultaneous expression of epithelial and mesenchymal marker genes.

### TGF-β1 induces invasive properties in CRC PDT1

Next, Reactome pathway analysis (32) was employed on significant differentially expressed genes between PDT1 cultured in basal medium with or without TGF-β1. Notably, we observed enrichment of pathways associated with ECM organization/degradation, elastic fiber formation, and cell adhesion (**Fig. 6A**). Within these pathways, several cell adhesion molecules, including integrins (*ITGB3,7,8*), latent TGF-β binding proteins (*LTBP1-4*), and *L1CAM-*associated molecules, as well as ECM associated proteins such as collagens, fibronectin, and metalloproteinases (*MMP14,17*), were among the top deregulated genes (**Fig. 6B**). The upregulation of the transmembrane protein *L1CAM*, which is highly expressed in metastasis initiating cells in CRC (33)*, MMP14* and *ITGA5*, which represent key molecules for ECM reorganization and elastic fiber formation, was additionally confirmed by qRT-PCR (**Fig. 6C**). Collectively, these findings suggest substantial alterations in the mechano-chemical properties of PDTs grown under TGF-β1 conditions, probably enhancing their invasive potential. Thus, Matrigel® trans-well invasion assays were performed to investigate the invasive properties of PDT1 under TGF-β1 exposure. While PDTs cultured in basal media exhibited no invasive potential, TGF-β1 treatment induced high invasive capacities of PDTs, represented by substantial migration of tumor cells through the pores of the trans-well mesh (**Fig. 6D**).

**Figure 6:**
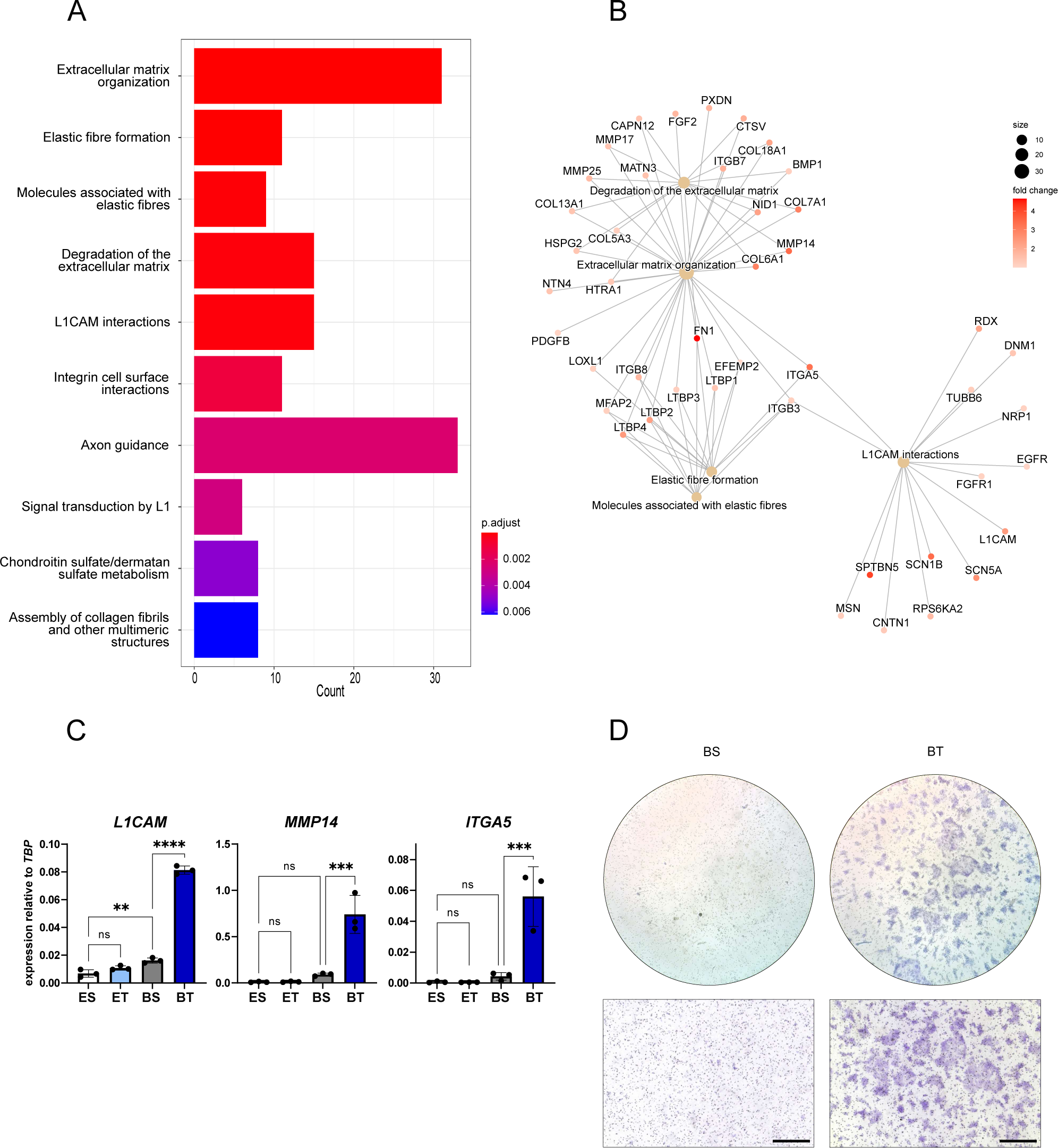
TGF-β1 induces invasive properties in PDT1. **A** Bar chart representing the top significantly enriched pathways in BT (basal medium + TGF-β1) *versus* BS (basal medium + solvent) conditions, based on Reactome pathway analysis of significantly upregulated genes (Padj < 0.05 and LFC > 1) from RNA-seq analysis between the two conditions (n=3). X-axis represents the counts of individual genes upregulated in the respective pathway. Color gradient indicates significance of the enriched pathway based on Padj. **B** CNET (Gene-Concept Network)-plot representing connections of genes among the five top-enriched Reactome pathways (according to Padj) from (A). Circle size indicates gene count for each pathway. Dot color for each gene indicates LFC change of the gene according to the color gradient. **C** qRT-PCR analysis of genes associated with invasion (*L1CAM*, *MMP14*, *ITGA5*) relative to the reference gene *TBP* of PDT1 cultured in different conditions (ES: ENAS + solvent; ET: ENAS + 5 ng/ml TGF-β1, BS: basal medium + solvent; BT: basal medium + 5 ng/ml TGF-β1). Graphs show the mean ± SD from 3 replicates (n=3). Statistical significance was calculated with GraphPad Prism version 8 using ordinary one-way ANOVA followed by Tukey’s multiple comparison test with 95% confidence interval: ns P > 0.05; *P f 0.05; **P f 0.01; ***P f 0.001; ****P f 0.0001. **D** Representative bright-field images of trans-well invasion assays in basal medium + solvent (BS) or in basal medium + 5 ng/ml TGF-β1 (BT) towards FCS as chemo-attractant. Invasive cells were visualized by crystal violet staining. Overview (top) and higher magnification image (bottom) are shown. Scale bar: 500 µm (lower figure).

In conclusion, TGF-β1 treatment significantly upregulates pathways involved in ECM remodeling and partial EMT characteristics of PDTs *in vitro*, thereby prominently enhancing their invasive properties.

## Discussion

Classical TGF-β studies utilized cell lines grown in 2D, which does not faithfully replicate several aspects important for EMT such as invasive properties, cell morphology and polarity, or cell-cell and cell-ECM interactions (34–36). The advent of innovative organoid and tumoroid models has provided suitable tools and novel insights into the effects of TGF-β on healthy and malignant cells (4,13,18,37,38). However, research on CRC tumoroids delivered partially inconclusive findings regarding the potential mechanisms underlying the TGF-β duality in CRC. Normal colon organoids, adenoma-derived organoids, as well as genetically engineered *BRAF*^V600E^ organoids, displayed phenotypic and transcriptional changes, indicating a critical importance of TGF-β already in precursor lesions to direct specific adenoma subtypes to the aggressive mesenchymal CMS4 CRC subtype (37). Studies on human tumoroids showed divergent results, demonstrating either a tumor-suppressive response and minimal changes in EMT marker gene expression (4) or *KRAS* dependent resistance to TGF-β treatment through downregulation of the pro-apoptotic protein BIM (13). Our study included five primary CRC PDTs with intact TGF-β signaling and different *KRAS* mutations. Notably, only one of the five lines, which harbored an atypical *KRAS*^Q22K^ mutation (PDT1), maintained proliferation and showed phenotypic changes. The *KRAS*^Q22K^ mutation occurs in about 0,2% of CRCs, and was previously associated with a 300-fold higher activity for inducing ERK2 *in vitro* compared to the frequently occurring exon 2 *KRAS*^G13D^ mutation (39). The remaining four primary PDTs harbored *KRAS*^G12D^ (PDT2,5), *KRAS*^G12V^ (PDT4) or exon 4 *KRAS*^A146T^ (PDT3) mutations and exhibited tumor-suppressive effects upon TGF-β1 treatment. Importantly, the different *KRAS* mutations have been associated with diverse treatment responses to chemotherapy and tyrosine kinase receptor targeting therapy, such as EGFR therapy, as well as patient outcome (40). Our data suggest that individual *KRAS* mutations result in different activation levels of MAPK signaling via pERK2, resulting in a disbalance of pro- and anti-apoptotic signaling via BIM and BCL-XL. Thus, the *KRAS* mutation status might be directly linked to TGF-β1 responses, impacting on tumor aggressiveness and therapy responses of the patients.

Organoids and tumoroids are classically cultured in media promoting their stem cell properties (41). However, since TGF-β signaling is highly context dependent, we here utilized an adapted medium, which only contained EGF as a growth factor and omitted classical ENAS medium factors and TGF-β pathway inhibitors for PDT cultivation. As expected, this resulted in downregulation of stem cell genes and upregulation of genes associated with specific colon cell subtypes including enterocytes and various secretory cell types. We previously observed similar phenotypes of PDTs upon co-culture with cancer associated fibroblasts (CAFs), which secrete several growth factors and cytokines, enabling the growth of tumoroids without the addition of niche factors (42). The observed diversity in colonic cell subtypes in this co-culture system resembled the *in vivo* tumors to a high degree (42). Therefore, we suggest media allowing for the differentiation of PDTs recapitulate *in vivo* tumor characteristics better than the commonly used ENAS medium and is thus more suitable to study TGF-β responses. The critical assessment of media conditions was also highlighted in a recent report on the drug response of PDTs derived from high-grade serous ovarian cancer (43).

Besides the upregulation of mesenchymal markers, the PDT1 line retained some epithelial characteristics following TGF-β1 treatment. This is reminiscent of partial EMT, which has been described for several cancer entities including breast, lung, and CRC, and is characterized by a heterogeneous population of tumor cells in different states of EMT (44). Notably, tumors displaying this incomplete acquisition of mesenchymal features possess the highest metastatic potential (44). Along these lines, we observed deregulated expression of genes and pathways involved in ECM organization/degradation, elastic fiber formation, and cell adhesion, which was paralleled by significantly increased invasive properties of tumoroid cells upon TGF-β1 stimulation. Moreover, some marker genes including the TF *PAX6* were specifically upregulated in basal medium following TGF-β1 treatment. An oncogenic role for PAX6 was previously reported for different cancer entities including CRC, breast, and non-small cell lung cancer (45–47). *PAX6* gene expression is regulated by canonical TGF-β1 signaling through SMAD3 binding to its promoter region (48). Moreover, PAX6 can interact with the MHC1 domain of different SMADs including SMAD3 and SMAD1 (49), suggesting that PAX6 represents a context-dependent transcription factor for CRC progression and metastasis, in line with the observed EMT phenotype. Together, these findings suggest essential tumor cell intrinsic capabilities to remodel the ECM and allow increased tumor cell motility, which was previously suggested to depend largely on stromal fibroblasts in the TME (50).

In conclusion, our findings show that context-dependent effects of TGF-β can be replicated *in vitro* in CRC PDT models, and most likely depend on the presence of aggressive *KRAS* mutations. Our findings underline the tumor cell-specific effects of TGF-β for the induction of a partial EMT state, matrix remodeling, and invasion, highlighting the value of PDTs for personalized medicine.

## Methods

### Statistics

Statistical evaluations were conducted using Prism software (GraphPad) version 8. The analysis incorporated a one-way analysis of variance (ANOVA), followed by Tukey’s multiple comparison test with 95% confidence interval and significance was determined at an (Padj) < 0.05.

### PDT cultivation

PDT models were established as described in (42). For cultivation, PDTs were maintained in droplets of Matrigel® in colon organoid medium as previously reported (41) without Wnt3a and R-spondin, referred to as ENAS medium (**supplementary table 1**) PDTs were routinely passaged at intervals of 7 days using TrypLE (Gibco, Cat. No. 12605010) and pipetting to dissociate the cells.

### Mutational profile of PDTs

After genomic DNA was isolated with QIAamp DNA Mini Kit (Qiagen, Cat. No. 51304), panel sequencing using the Ion AmpliSeq™ Cancer Hotspot Panel v2 (Thermo Fisher, Cat. No. 4475346) was done to infer driver mutations of individual lines. In addition, whole exome sequencing was performed for PDT8 (MSI-high) to identify mutations in *TGFBR2*.

### Embedding of PDTs and Immunohistochemistry

PDTs were fixed with 4,5% paraformaldehyde (PFA) and dissolved into agarose domes, which were dehydrated and embedded in paraffin (details of this procedure can be found in Supplementary information, SI). For Hematoxylin and eosin (H&E) staining 2-3 µm thick sections of embedded PDTs or corresponding tumor tissues were used.

### TGF-β1 treatment of PDTs

Single cells were seeded in a concentration of 2000 cells/ 10 µl in 50% Matrigel/ PBS domes. After 72 hours in ENAS + ROCK inhibitor, medium was replaced to different conditions: ENAS + solvent (0,2 mM HCL/PBS + BSA), ENAS + 5 ng/ml TGF-β1 (Recombinant Human TGF-β1 Protein, R&D Systems, Cat. No. 240-B). Basal medium + solvent. Basal medium + 5 ng/ml TGF-β1. PDTs were treated for a time course of 10 days and medium was changed every other day. Detailed medium composition is provided in SI, supplementary table 1.

### Viability assays

Single cells were seeded in a concentration of 2000 cells/ 10 µl, as 8 µl 50% Matrigel/PBS domes in white 96 well plates as technical triplicates. Treatment was carried out as stated in TGF-β1 treatment. CellTiter-Glo 3D Cell Viability Assay (Promega, Cat. No. G9681) was used to detect viability. For this, medium was removed and 75 µl fresh BASAL medium without EGF and 75 µl CellTiter-Glo 3D reagent was added. Measurement was carried out according to manufacturer’s instructions using plate reader synergy H1 (Bio Tek Agilent). The following formula was used to calculate well viability: well viability= ((well value - average positive control) / (average vehicle control-average positive control)) *100%, whereby positive control refers to Staurosporin 5 µM control (Sigma Aldrich, Cat. No. 569397), and the vehicle control refers to ENAS + solvent. Values were plotted in GraphPad Prism version 8.

### RNA isolation and gene expression analysis

RNA was isolated directly after treatment with RNeasy Kit (Qiagen, Cat. No. 74104) using the on-column DNA digestion protocol following the manufacturer’s instructions. For qRT-PCR, iScript™ cDNA Synthesis Kit (Bio Rad, Cat. No. 1708891) was used for cDNA preparation and Luna® Universal qPCR Master Mix (NEB, Cat. No. M3003L) was used for subsequent qPCR. Relative gene expression levels were calculated with the delta CT method using TATA-Box Binding Protein *(TBP)* as a reference gene (Primers are listed in SI supplementary table 2). Each sample was measured in technical triplicates. Isolated RNA was processed for mRNA-sequencing, all details of sequencing and gene expression analysis can be found in SI.

### Immunofluorescence staining

Single cells were seeded into 8-well glass bottom slides (IBIDI, Cat. No. 80827). Following treatment as stated in TGF-β1 treatment section, PDTs were fixed and stained directly in these slides, detailed procedure and list of used antibodies can be found in SI and supplementary table 3.

### Protein isolation and Western blotting

To collect PDTs, the culture medium was removed, the domes were rinsed with PBS and resuspended in 350 µl Cell Recovery Solution (Corning™, Cat. No. CLS354253). The mixture was incubated for 45 minutes at 4°C, then washed with cold PBS. PDTs were harvested by centrifugation. Post-collection, the pellets were snap-frozen and subsequently stored at −80°C. Protein isolation and western blotting was carried out as described earlier (51), with minor deviations: 10 µg Protein was used for each blot and BSA was used for blocking of membranes. Antibodies used are provided in SI supplementary table 3.

### Trans-well invasion assay

35 000 single cells were seeded in 50 µl 10% Matrigel/PBS into Matrigel coated trans well plates (Corning, Cat. No. 354480). After 72 hours in ENAS + ROCK inhibitor, PDTs were cultured for five days in BASAL medium + solvent or 5 ng/ml TGF-β1. In lower wells BASAL medium was added. After this pre-treatment, PDTs were treated further in these conditions, and in the lower well 300 µl BASAL medium + 10% FCS as chemo attractant was added. As negative control (no attraction) BASAL medium only was added. After 3 days medium (+/- TGF-β1) was changed in upper wells, and in lower wells 300 µl of medium with or without FCS was added. After five days of chemoattraction cells which migrated through the membrane and were attached at the lower side of the mesh, were fixed with 100% methanol, and stained with crystal violet (Sigma Aldrich, Cat. No. C6158).

## Data availability

The analysis of the RNA sequencing data is available within the results section and the supplementary information files. The RNA sequencing raw data will be publicly available in Gene Expression Omnibus, whereby PDT1 refers to CRC3.

## Acknowledgments

We thank: Helga Schacher, Sabrina Wohlhaupter, Astrid Haase for technical support, Tanja Limberger for providing protocols, Gabriel Wasinger for pathological support, Philipp Hofer and the Biobank team of the Department of Pathology for their support in sample storage. Wolfgang Mikulits and Helmut Dolznig for their critical discussion. Next-generation sequencing and initial data analysis was performed by the Biomedical Sequencing Facility at CeMM, Research Center for Molecular Medicine of the Austrian Academy of Sciences.

## Contributions

**GE** acquired funding and supervised the project. **TM** and **GE** designed the project and wrote the manuscript. **TM, PK**, **MM, PMS** performed and analyzed experiments. **LT** established PDT cultures and protocols. **TM, CUPM, KD** and **RST** performed data analysis and visualization. **JP, JuK, VSA** provided PDT lines. **MB** provided patient material and clinical data. **JeK** helped with writing and scientific discussion. **AT** performed pathological assessment. **LWO** and **LM** carried out PDT panel sequencing. All authors reviewed and approved of the manuscript.

## Ethics declaration

All experiments performed were adhering to the “Guidelines for Good Scientific Practice” and in alignment with the most recent “Declaration of Helsinki.” Patient material was used upon signed consent, only. The ethics governing this research received approval from the IRB of the Medical University of Vienna (N1248/2015).

## Consent for publication

All the authors approved the final version of the manuscript and agreed to its publication.

